# Neuronal activity triggers uptake of hematopoietic extracellular vesicles *in vivo*

**DOI:** 10.1101/735670

**Authors:** Ivan-Maximiliano Kur, Pierre-Hugues Prouvot, Ting Fu, Wei Fan, Felicia Müller-Braun, Avash Das, Saumya Das, Thomas Deller, Jochen Roeper, Albrecht Stroh, Stefan Momma

**Author notes:** Lead Contact: All related correspondence should be addressed to Dr. Stefan Momma, Institute of Neurology (Edinger Institute), Frankfurt University Medical School, D-60528 Frankfurt, Germany.

## Abstract

Communication with the hematopoietic system is a vital component of regulating brain function in health and disease. Traditionally, the major routes considered for this neuroimmune communication are either by individual molecules such as cytokines carried by blood, by neural transmission, or in more severe pathologies, by entry of peripheral immune cells into the brain. In addition, functional mRNA from peripheral blood can be directly transferred to neurons via extracellular vesicles (EVs) but the parameters that determine their uptake are unknown. We show that transfer of EVs from blood is triggered by neuronal activity *in vivo*. Importantly, this transfer occurs not only in pathological stimulation but also by neuronal activation caused by the physiological stimulus of novel object recognition. This discovery suggests a continuous role of EVs under pathological conditions as well as during routine cognitive tasks in the healthy brain.

## Introduction

Extracellular vesicles (EV) emerge as ubiquitous signaling agents that can transfer functional proteins, nucleic acids and lipids between cells *in vitro* and *in vivo* [1–4]. The ability to transfer such a range of different molecules is particularly intriguing in the communication between the brain and the periphery. EVs can contain signature molecules that may reflect the physiological state of secreting neural cells, passing the blood-brain barrier and thus making them obvious targets for diagnostic use. Vice versa, EVs may be a valuable tool for the delivery of therapeutic molecules to the brain [5,6]. However, the physiological parameters that regulate EV signaling to the brain *in vivo* are poorly understood. Here we investigate if the uptake of EVs released by peripheral blood cells is triggered by neuronal activity. In order to track EV signaling *in vivo*, we used a transgenic mouse model developed by us that can be leveraged to trace the transfer of functional mRNA by blood-derived EVs *in vivo* using the Cre-LoxP recombinase system[2,3]. Starting from the observation that peripheral inflammation induced by injection of LPS leads to a massive onset of blood to brain signaling via EVs, we made similar observations in models that are more or exclusively characterized by neuronal activation such as kainate (KA) injection, pharmacological inhibition of the proteasome in dopaminergic neurons and, most specifically, by optogenetic induction of neuronal activity. Furthermore, neuronal activity and concomitant EV uptake can also be induced by behavioral cues as in novel object placement *in vivo*. Together, this argues for a continuous communication via transfer of functional molecules contained in EVs in various pathological and physiological states. The transfer of molecules by EVs that were so far not considered in intercellular signaling may be of great potential importance[1].

## Results

### LPS induced peripheral inflammation leads to widespread EV uptake in the brain

The transgenic mouse model that we were using for our study relies on the fact that Cre mRNA is expressed in hematopoietic cells under the vav1 promoter [7] and sorted into EVs that are then released into the bloodstream. The exact mechanism of how the Cre mRNA is sorted into exosomes is unclear but it is probably a simple reflection of high cellular expression [8]. Upon entering a target cell, Cre mRNA is translated to functional protein leading to the irreversible onset of a marker gene expression, here EYFP, mediated by Cre recombinase activity (Fig. 1A). In the healthy animal, no or very few recombined neural cells indicative of EV signaling are present in the brain[2,9] (Fig 1B). However, after peripheral inflammation by two daily, i.p. injections of LPS (1mg/Kg), we observe frequent recombination events in the hippocampus (HC) (Fig 1C-G and Fig 1J) and substantia nigra (SN) (Fig 1H to J) but also in other brain regions (Supplemental Fig 1). For the dentate gyrus (DG) of the HC, marker gene NeuN-positive neurons reached on average 12.3% (±4.7% SD, n=5), and Iba1-positive microglia 52.3% (±3% SD, n=5). In addition, we observed 21.5% (±4.4% SD, n=5) of TH-positive dopaminergic neurons (DA) expressing EYFP and 45.1% of Iba1-positive microglia (±6.7% SD, n=5) in the SN (Fig 1H to J). For the SN, we could also frequently observe recombined cells with neuronal morphology that were TH-negative (Fig 1H). Given the prominent role of inflammation in Parkinsons disease, this may point to a previously unidentified factor linking these events. We very rarely or never observed cells of endothelial (CD31), astrocytic (GFAP) or oligondedroglial (Olig2) lineages co-expressing the marker gene. As a complement to our vav-iCre mice and in order to gain further insight into the role of the origin of EVs in the selectivity of cell targets, we analyzed brains from EpoR-iCre mice after peripheral LPS injection. In EpoR-iCre mice, iCre is expressed under the control of the erythropoietin receptor, restricting its expression to the erythroid lineage[10]. Analysis of brain sections revealed an absence of neuronal recombination while still observing recombined Iba1-positive microglia (Fig. 1K and L) indicating that the targeting of EVs to specific cells is determined, at least in part, by their cell of origin.

**Figure 1.**
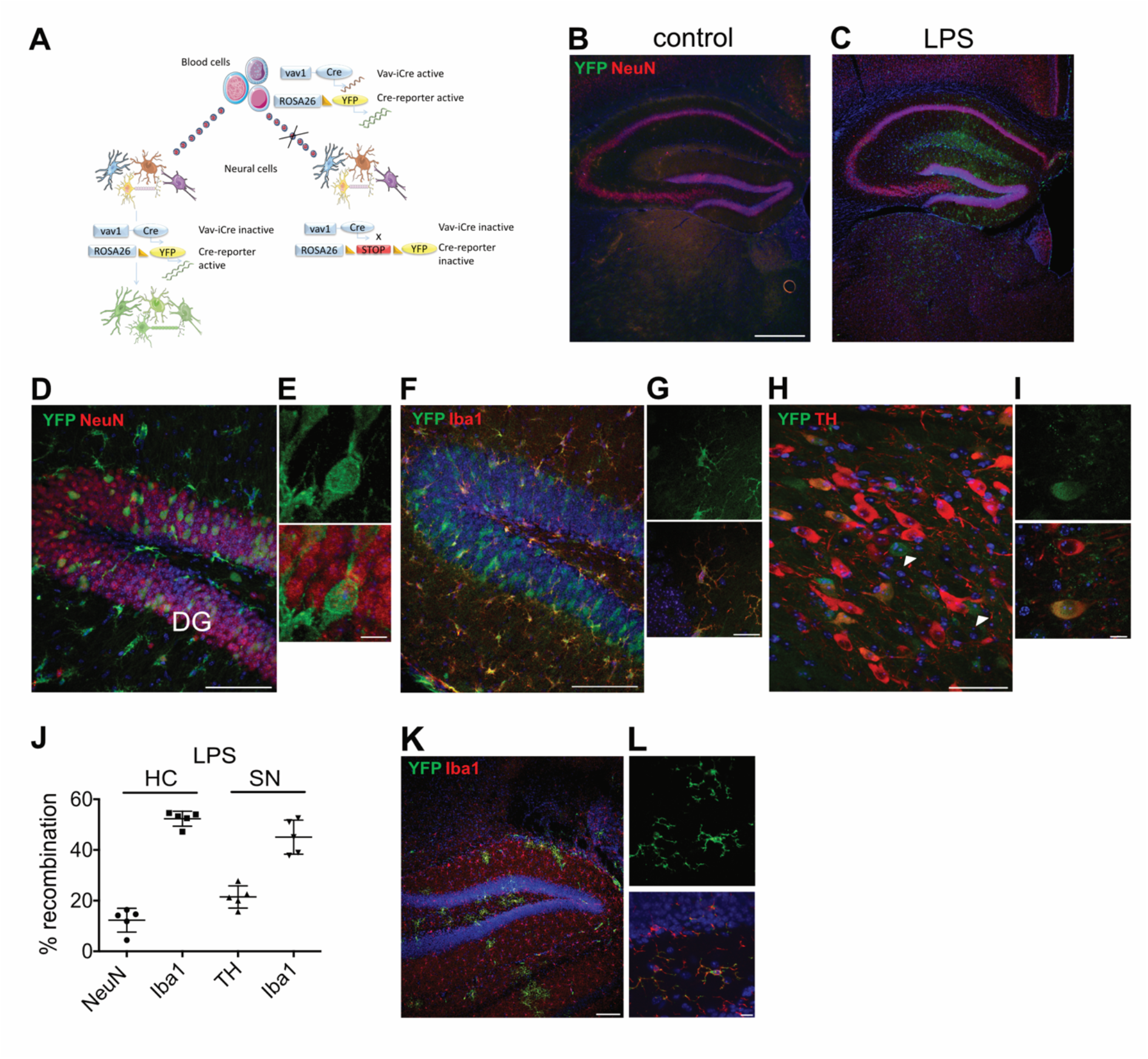
Peripheral stimulation initiates widespread EV uptake in the brain. (A) Schematic representation of the method to map EV mRNA transfer from blood to brain. Cre mRNA contained in blood derived EVs is taken up in neural cells, leading to excision of the stop-loxP site and induction of marker gene expression. (B) Hippocampus from a control vav-iCre-R26EYFP mouse compared to 48h after i.p. LPS injection, showing frequent recombination events in the hippocampal area (C-G). Recombined cells are predominantly neurons and microglia (D-G), including dopaminergic neurons in the substantia nigra (H+I). Both structures show similar levels of recombination (J). (K+L) Recombination after LPS treatment in EpoR-iCre mice is restricted to microglia. Scale bars 500µm in B; 100µm in D, F; 10µm in E, G, I, L; 50µm in H.

### Kainic acid induced neuronal activity

To probe for a more specific stimulation of neuronal activity we injected KA, a neuroexcitatory agonist for kainate receptors that is frequently used as an epilepsy-like model. KA injection (single i.p. injection of 10mg/Kg) lead to the transfer of Cre mRNA to the brain, indicated by marker gene expression in neurons as well as microglia (Fig 2A to C). Recombination levels in DA neurons (Fig 2D and E) were similar to those seen in DG granule neurons in the HC but much lower in microglia compared to the LPS model. Altogether, levels of marker gene expression were comparable to those after LPS injection (NeuN: 18.5% ±14.6% SD, n=5) although with a higher variability, as well as a shift to more neuronal-and less microglial recombination in the hippocampus (Iba1: 35.2% ±16.3% SD, n=5) (Fig 2F). Variations in the SN were less pronounced (S3 Fig) (TH: 23.2% ±3.1% SD; Iba1 29.2% ±7.1%, n=5).

**Figure 2.**
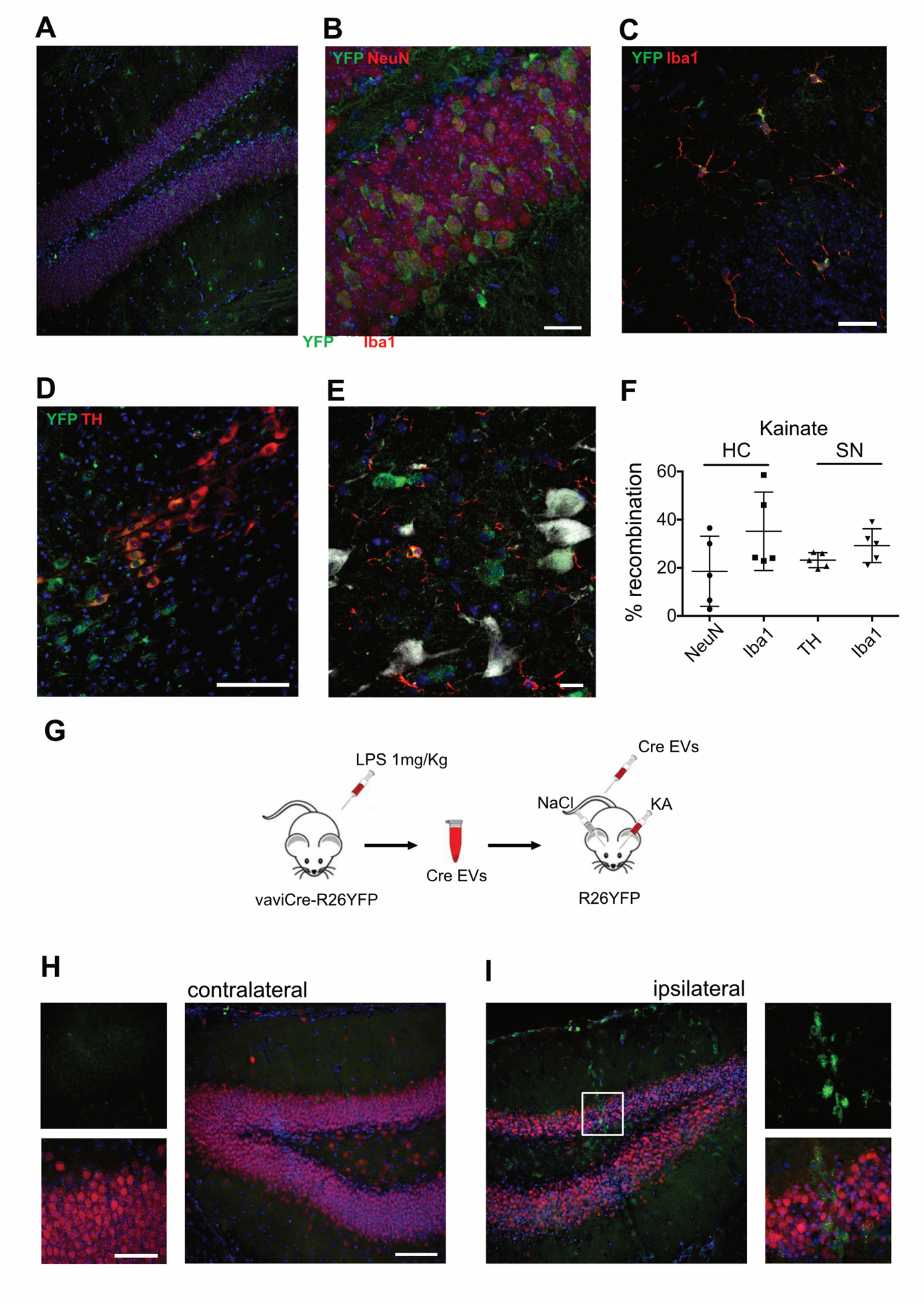
Recombination in HC and SN after Kainate injection. (A) Hippocampus with DG showing multiple recombination in neuronal and non-neuronal cells. (B) Magnified view of DG with marker gene-expressing granule neurons as well as Iba1-positive microglia in (C). (D) SN with TH-positive but also TH-negative neurons expressing EYFP as well as microglia in (E). (F) Quantification of marker gene-positive cells after KA injection. (G) Experimental scheme for peripheral EV injection followed by local neuronal activation by intracerebral injection of KA with saline solution into the contralateral hemisphere as control. (H) In the control hemisphere, no EYFP-positive cells are visible except meningeal macrophages on the brain surface. (I) In contrast, in the locally stimulated hemispheres, marker gene-positive neural cells can be observed in the DG. Scale bars 100µm A, D, H right panel; 10µm E; 20µm B; 50µm H right panel.

Next we wanted to control for the possibility of immune cell infiltration from the periphery caused by the injection of LPS or KA. Peripheral macrophages can be distinguished from brain resident microglia by their expression of the integrin subunit alpha 4 (Itga4/Cd49d) [11]. Immunohistochemical staining of brain sections from both conditions (n=3 for each) lead to no observable CD49d-positive cells in the brain parenchyma thereby ruling out this possibility (Supplemental Fig 2).

The presence of both the Cre expressing and Cre reporter construct in the same cells leaves the theoretical possibility of marker gene induction being caused by unspecific expression of Cre recombinase. To address this issue, we injected KA into the hippocampus of one hemisphere of ROSA26-EYFP Cre-reporter mice, lacking endogenous Cre expression, and saline solution into the contralateral side. Subsequently, we injected EVs isolated from the plasma of LPS stimulated vaviCre mice into the tail vein (Fig 2G). Analysis of brains after 48h showed recombination events only at the site of KA injection, but not in the contralateral hemisphere (Fig 2 H and I). Thus, recombination could have only been induced by EVs entering the brain via the peripheral circulation. Furthermore, local neuronal activation by KA injection was necessary and sufficient for EV uptake whereas injection of carrier solution alone did not lead to any EV uptake. Overall, we interpreted our results as an indication that neuronal activity may be a factor regulating EV uptake and turned to experimental paradigms suitable to specifically address this issue.

### Inhibition of ubiquitin-proteasome system leads to specific EV uptake in DA neurons

Transient increases in neuronal firing are a common pathological feature of Parkinson’s disease and well described in animal models where protein degradation via the ubiquitin-proteasome system is impaired either genetically or pharmacologically[12,13]. We unilaterally infused the selective proteasome inhibitor, epoxomicin[14] into the ventral midbrain as described previously[13] (Fig 3A). Two weeks after infusion, mice were killed and the brains were processed for quantitation of EYFP-expressing DA neurons in the SN. We observed no recombination events in TH-positive neurons in the contralateral site of infusion and no recombination in animals that received an infusion of carrier solution (n=3) (Fig 3B). In the ipsilateral site of infusion, we could detect EYFP-TH double positive neurons (Fig 3C to E) (8.6% ± 3.1% SD, n=4). These results are in line with a highly specific uptake of EVs in DA neurons when firing frequencies in DA SN neurons are increased. Interestingly, we could detect none to only very few recombined microglia which may be due to the reported anti-inflammatory properties of epoxomicin[14].

**Figure 3.**
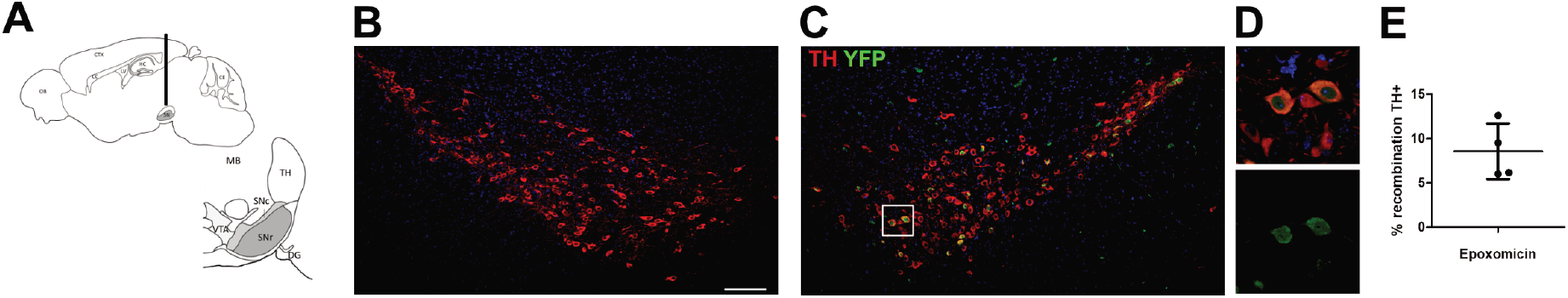
Neuronal activation is sufficient to trigger EV uptake. (A) Unilateral infusion of the selective proteasome inhibitor epoxomicin into the ventral midbrain leads to increased *in vivo* firing frequencies of DA SN neurons. Two weeks after infusion no marker gene expressing DA neurons were discernible in the contralateral hemisphere (B). In contrast, DA neurons in the ipsilateral hemisphere frequently took up EVs leading to marker gene expression (C-E). Scale bar, 100µm in B.

### Specific induction of neuronal activity by optogenetic methods

To directly stimulate neuronal activity with high specificity, we turned to an optogenetic approach[12]. Channelrhodopsin-2 (ChR2) is a rapidly gated blue light-sensitive cation channel leading to membrane depolarization that can evoke action potential firing[15] in neurons. We injected Adeno-Associated Virus (AAV) encoding ChR2 fused with mCherry under an excitatory-neuron-specific CamKIIA promoter into the primary visual cortex of vav-iCre-ROSA26-EYFP mice (experimental scheme in Fig 4A). We first assessed the expression profile of the opsin ChR2 fused with the fluorophore mCherry. We found strong and membrane-bound expression of ChR2-mCherry mainly in layer V, and to a lesser extent in layers II and III (Fig 4B and C). We next tested the functionality of our optogenetic approach by conducting *in vivo* electrophysiological field potential recordings. For that, we illuminated the opsin-expressing region with an optic fiber emitting blue light of varying light intensities, 4 weeks after rAAV2-CaMKIIa-hChR2-(H134R)-mCherry-WPRE-pA virus injection. We found a typical sigmoidal dose-response curve [16] (Fig 4D and E). Based on these data, we chose the light intensity for subsequent experiments, which reliably evoked strong neuronal responses. To control for unspecific induction of EV uptake by the experimental procedure, we also analyzed brains from animals that received an AAV injection (n=4) without optical stimulation (Fig 4F), or where we performed an optical stimulation (n=4) without prior injection of AAV (Fig 4G). In both cases, the treatment did not induce any neuronal recombination events. In contrast, in all animals with a combination of an AAV injection and optical stimulation (n=4), we observed recombined, EYFP- and NeuN-positive neurons (Fig 4H). The number of marker gene-positive cells in the area of ChR2-mCherry expression indicated by the white box in Fig 4H was 44.6 cells per 0.01 mm^3^ (± 6.6 cells per 0.01 mm^3^ SD) with 3.7% (±3.5% SD) of EYFP-positive cells being microglia. However, only a part of recombined neurons co-expressed ChR2-mCherry and most recombination events were actually outside the injection area in adjacent cortical areas and in the hippocampus (Fig 4H to K). Indeed, virally transduced neurons send projections to the contralateral hemisphere with accompanying recombination in the surrounding area (Fig 4L and M). We interpret this as a network stimulation effect whereby connected neurons will increase firing thus leading to uptake of EVs in cells that were not directly stimulated. We did not observe any recombination events in areas distant from the stimulation site such as the forebrain and cerebellum (Fig 4N and O) underlining the highly localized stimulation.

**Figure 4.**
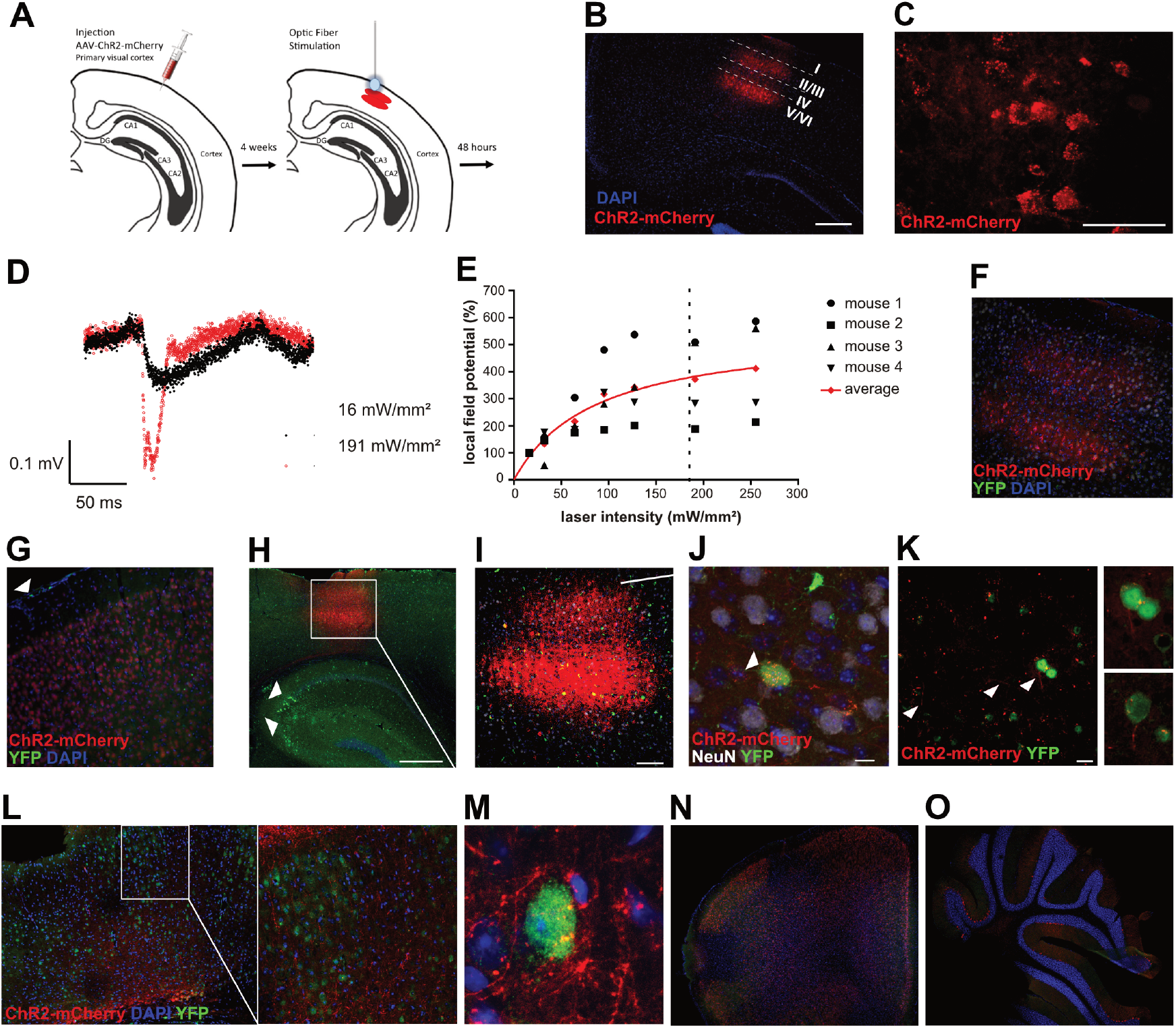
Neuronal activation by optogenetic stimulation. (A) Scheme for the injection of ChR2-mCherry-AAV in layer 5 of the primary visual cortex and optical stimulation. (B) Expression pattern of ChR2-mCherry in primary visual cortex (V1), 4 weeks after AAV injection. Scale bar: 500 µm. Layer-specific expression of ChR2-mCherry, mainly in layer V of V1. (C) Membrane-bound expression of ChR2-mCherry Scale bar: 50 µm. (D) Representative *in vivo* local field potential (LFP) traces from mouse 4 upon optic-fiber-based illumination with blue light with an intensity of 16 and 191 mW/mm^2^ at the tip of the fiber. (E) Normalized LFP amplitudes, plotted against light intensity. Averaged amplitude was fit into a nonlinear curve. Dotted line marked the light intensity used in subsequent experiments (~185 mW/mm^2^). (F) Control sections of AAV injected mice without optical stimulation as well as from mice that received only an optical stimulation without AAV injection (G)(white arrowheads indicate marker gene positive meningeal macrophages). (H) Recombination occurs in ChR2-mCherry-positive and -negative neurons but also in the hippocampus (arrowheads) (H-K). Projections of ChR2 expressing neurons extend to the contralateral side and recombination can also be observed in target areas (L-M). No marker gene-positive cells can be observed in sections from the forebrain (N), or the cerebellum (O). Scale bars 500µm in B+H; 100µm in I; 10µm in J; 20µm in K; 1mm in N.

### Hippocampal neuronal activity induced by novel object placement

So far, in our experiments we elicited the induction of EV uptake via pathological events or ectopic stimuli. Now we probed whether increased neuronal activity induced by behavioral paradigms would suffice for neuronal EV uptake. To this end, we chose the induction of hippocampal activity by environmental novelty. The hippocampus is involved in the formation of contextual memory and it has been shown that addition of novel objects into a mouse cage leads to a selective increase in neuronal c-fos expression in the hippocampus[17]. We placed several new objects into a cage with vav-iCre-ROSA26-EYFP mice. The objects were removed again after one hour (Fig 5A). After 48h animals were killed and their brains analyzed at different rostro-caudal levels. We could not detect marker gene-positive neurons in animals that were housed in cages that were left unchanged (n=4) (Fig 5B). In the animals from the cages with new objects (n=9) we observed EYFP-positive neurons in the hippocampus (Fig 5C to E) in all but one animal. We counted marker gene-positive cells at a frequency of 117 cells per 0.01 mm^3^ (±14.1 cells per 0.01 mm^3^ SD, n=3) for the DG and 45 cells per 0.01 mm^3^ (±13.8 cells per 0.01 mm^3^ SD, n=3) for the CA1 and CA2 hippocampal subfields with 13.7 % (±2.1 SD, n=3) of EYFP-positive cells being microglia (Fig 5F). We did not, or only very rarely, detect any EYFP-positive neurons in the forebrain, hindbrain or cerebellum where no activation would be expected (Fig 5G to I).

**Figure 5.**
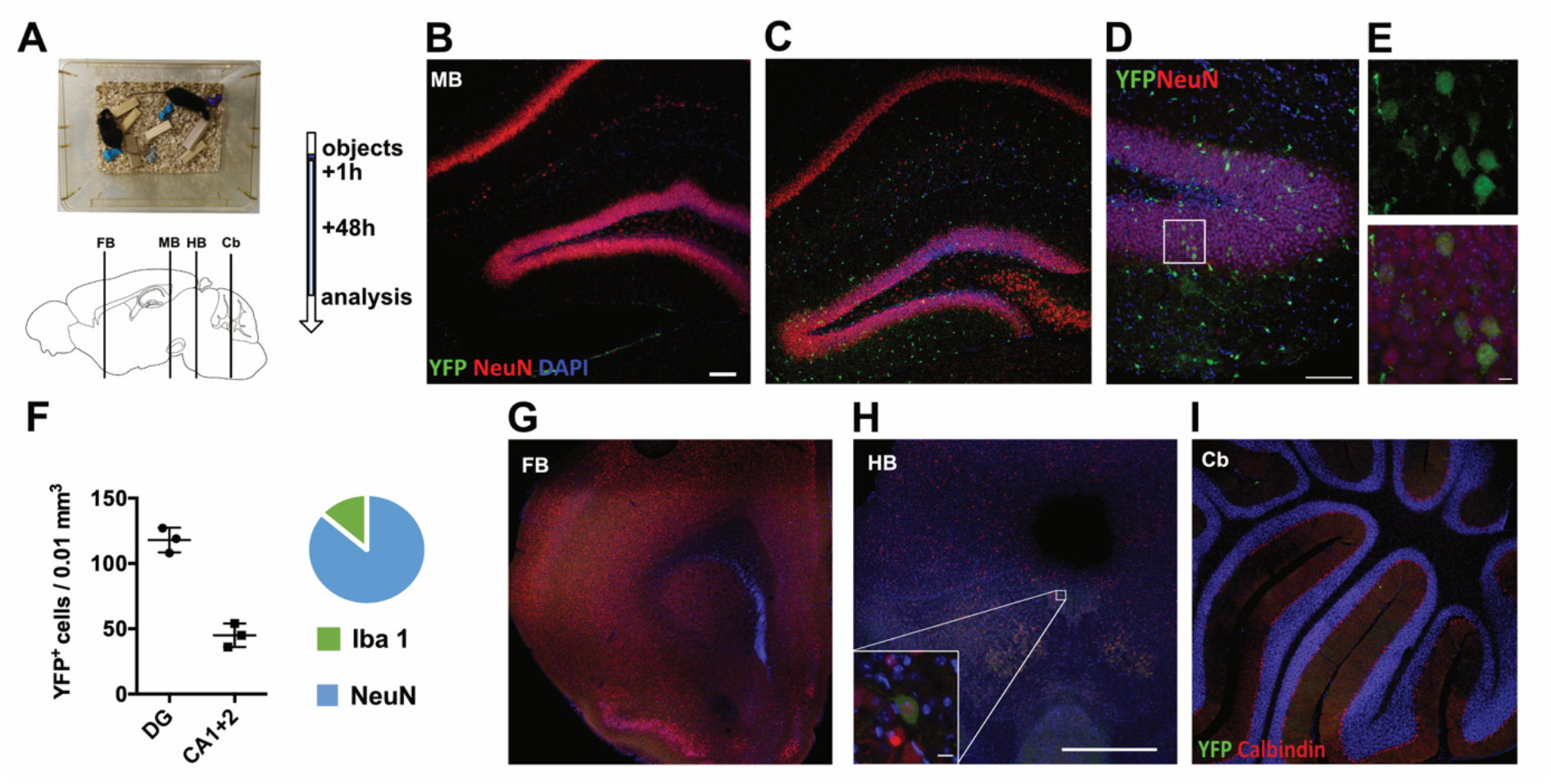
Neuronal stimulation by novel object placement. (A) Novel objects were placed in mouse cages and left for one hour. Brains were analyzed along their entire rostro-caudal length 48h thereafter. (B) No marker gene expression was observed in controls in contrast to placement mice (C). Recombination occurred mainly in neurons in hippocampal areas DG and CA1 and CA2 but also in some microglia (D-F). Marker gene expression was absent or very rare in other brain areas such as forebrain (G), hindbrain (H) and cerebellum (I). A single marker gene-positive neuron in the hindbrain is shown in insert (H). Scale bars 100µm in C, E; 500µm in H, 10µm in F and insert in H.

## Discussion

In summary, we demonstrate that functional transfer of molecules contained in EVs (Extracellular vesicles) that are released from hematopoietic cells and taken up by neurons is widespread and consistent with the notion that EV uptake is triggered by neuronal activity both under pathological and physiological conditions.

Conceptually, in the experimental settings we used, EV signaling appears to follow more a “demand-pull” rather than a “supply-push” principle in that the status of the target cell determines uptake. The blood-brain-barrier does not seem to be a major factor in limiting EV passage into the brain which was surprising given that the conditions we used are not obviously damaging the BBB and since we never observed infiltrating peripheral blood cells. A possible explanation is that neuronal activity which leads to the uptake of EVs might simultaneously regulate BBB permeability as shown previously [18]. This would present an interesting activity-dependent neurovascular coupling affecting neuronal regulation by peripheral factors.

For neurodegenerative diseases like PD, research in the EV field focuses mainly on the intercellular transfer of misfolded proteins or on the use of EVs as diagnostic markers for neurodegenerative processes[19,20]. Our results now suggest an ongoing role of EVs that may impact disease onset and progression. We do not know yet whether EV uptake is beneficial or detrimental for neurons, and whether the cell of origin determines its function. Notably, oligodendrocyte-derived EVs can have trophic effects on neurons *in vitro*[21] and macrophage-derived exosomes, containing functional NADPH oxidase 2 complexes promote axonal regeneration [22]. On the other hand, neuronal activity has been linked to tau propagation[23] and uptake of EVs may at least in part explain this transneuronal spread.

The surprising fact that we observe EV uptake under physiological conditions, induced by neuronal activity due to an enriched environment, suggests a role of EV signaling going beyond a reaction to pathology-induced cellular changes. This opens the question if cognitive processes or behavior can be influenced by EVs originating from the outside of the central nervous system in the medium or long term. We have to consider that our transgenic animal model reports EV transfer from peripheral blood only, while EVs released from any other organ, including the entire microbiome, go unreported. Thus, the extent of an influence of EVs on cognitive processes in the brain may be substantial. Lastly, EVs show great promise in delivering drugs or even RNA across the blood brain barrier and efforts are being made to target these to specific cellular populations. Our results show that, at least in the case for neurons, EVs seem to naturally target cells that are stimulated both by pathological and physiological processes with high specificity.

## Acknowledgements

We thank Jadranka Macas, Maika Dunst, Tatjana Starzetz and Sonja Thom for technical support.

## Author contributions

IMK, TF and WF performed experiments and analyzed data. SD, AD and FMB designed and performed experiments, JR and TD designed experiments and contributed to methodology and resources. AS and PHP designed, performed and analyzed experiments, SM conceived the project, designed and performed experiments, analyzed data and wrote the manuscript.

## Materials and Methods

### Animals

All animal experiments and protocols were approved by the regional Ethical Commission for Animal Experimentation of the state of Hessen, Germany (ethical permissions Gen. Nr. F90/10, F94/12, FK/1090, F94/19), the state of Rhineland-Palatina (ethical permission G 14-1-040) and the Institutional Animal Care and Use Committee for Massachusetts General Hospital

All mice were group housed (maximum four mice per cage) under standard laboratory conditions with a 12:12h light:dark cycle with food and water provided ad libitum. All experiments were performed on 6-to 30-week-old mice during the day cycle. Cage enrichment experiments were conducted between 10:00 and 13:00 during the day cycle. Vav-iCre mice were a gift by Dimitris Kioussis[7] (JAX-mice stock number 008610). The vav-iCre line was cross-bred with the Cre reporter mice ROSA26-EYFP (JAX-mice stock number 005130). For the detection of double-transgenic offspring, a drop of blood was collected from each pup and processed for analysis by Fluorescence-Activated Cell Sorting (BD FACS Canto^TM^ II Flow Cytometer) for EYFP expression. The EpoR-iCre mice were obtained from Dr. Stuart Orkin (Boston Children’s Hospital) and are deposited in the Jackson Laboratories; these were cross bred with the ROSA26 mTom/mGFP mice for the experiments.

### Extracellular vesicle preparation

Peripheral blood from deeply anesthetized reporter or double-transgenic mouse previously injected with LPS was collected during perfusion. Platelet-free plasma was isolated through centrifugation steps (Heraeus Labofuge 400R) 2500 rpm 15min RT twice and diluted 1:1 with 10mM HEPES buffer (A6916,0125 AppliChem Panreac) in Millipore water. The solution was 0,45um filtered to remove larger particles and mixed with Polyethylene glycol (PEG) 8000 (Rotipuran, 0263.1, Roth) solution 1:5 for precipitating vesicles during an overnight incubation at 4°C. Samples were centrifuged at 1500g 30min 4°C and pellets were resuspended in 10mM HEPES buffer for an ultracentrifugation step (Sorvall WX Ultra Series 80, Thermo Fischer) at 100.000g for 90min. The pellets were resuspended in 10mM HEPES buffer and kept at 4°C for immediate use or at −80°C for longer storage.

### *In vivo* experiments – Stereotactic surgeries

For experiments using stereotactic surgeries for viral injections and optic fibers, mice were anesthetized with isoflurane (Floren, Abbott), 2% v/v in O_2_ and placed in a stereotaxic apparatus (Model 900 / 940, KOPF Instruments) with adapted components to allow mouse inhalation anaesthesia. Before surgery, Xylocain (AstraZeneca) was administered as a local analgesic at the incision site on the skin. For closing incisions, few drops of tissue adhesive Surgibond (190740, SMI sutures, Praxisdienst) were used.

For optogenetics assay, Adeno-associated virus (AAV, titer of 10^13^ genomic copies per ml) with CamKII promoter driven hChR2(H134R)-Cherry (VB4411, Vector Biolabs) was used. Intracranial injection was performed unilaterally with elongated glass capillaries (612-2401, Hirschmann, VWR) in a 35º angle and the injected volume was 1 µl (coordinates in millimetres from bregma M/L=2,5; A/P=-3,1; D/V=0,5). At the end of the infusion, needles were kept at the site for 4 min and then slowly withdrawn. Viral expression was assessed 4 weeks after surgery.

For light stimulation a 200 μm multimode fiber with a numerical aperture of 0.39 (Thorlabs, Munich, Germany) was positioned unilaterally (touching the cortex), over the site of the AAV injection. The fiber was coupled to a Solid state 488nm 50mW Laser (Lighthub, Rodgau-Dudenhofen Germany) controlled by the omicron proprietary software. The laser output power was adjusted to read 5,8 mW measured at the fiber (~185 mW/mm²). The laser was pulsed at 10Hz with 10ms pulse width at 0.1 Hz for 1h, using an external pulse stimulator (CED 1401 Cambridge Electronic Design, Cambridge, UK). After the stereotaxic surgery, the animals were left for two days to allow for expression of EYFP and then perfused for further analysis. Local field potential (LFP) recordings *in vivo* were performed using glass pipettes (impedance ~1 MΩ) in layer V of primary visual cortex 4 weeks after rAAV2-CaMKIIa-hChR2-(H134R)-mCherry-WPRE-pA virus injection. Mice were anesthetized with isoflurane by inhalation and placed on a warming pad (37°C). A silver wire was inserted into the cerebellum as the ground electrode. To induce neuronal activity, single blue light pulses pulse (10 ms) were administered at 0.1 Hz via a 200 μm-diameter multimode fiber. The laser output power at the fiber tip was adjusted to 0.5, 1, 2, 3, 4, 6 and 8 mW (corresponding to 16, 32, 64, 95, 127, 191 and 255 mW/mm^2^ at fiber tip). At each intensity, 10 sweeps were recorded and averaged. In Figure 3, LFP amplitudes were normalized by the amplitudes evoked at 16 mW/mm^2^ intensity.

To induce peripheral inflammation, 200µl of lipopolysaccharide (LPS from E. coli O55:B5, 1mg/ml/kg, L2637 Sigma) was injected intraperitoneal every 24h for 2 days (two injections in total). After the last injection mice were kept for 48h, then animals were perfused for further analysis.

For Kainic acid (KA) injection, 100ul of KA (2,5mg/ml, ab120100, Abcam chemicals) was injected intraperitoneal once. After 48h mice were killed and transcardially perfused for further analysis.

For epoxomicin infusion, craniotomy was performed at the following coordinates in millimetres from Bregma M/L=0,9; A/P=-2,85; D/V=4 and 10uM epoxomicin in 1% DMSO or 1%DMSO alone as a control were infused unilaterally using a micro-pump (UMP3-1, WPI, Berlin, Germany; 10-lL nanofil syringe, 33-gauge steel needle; flow rate of 100 nL/min). After a 2-week post-infusion period, the anesthetized animals were transcardially perfused for further analysis.

For the combination of i.v. EV injection and local neuronal stimulation, ROSA26-EYFP mice received an intracranial injection of 0.5 µg of KA in 1 µl of saline solution (ipsilateral) and 1 µl of saline solution only (contralateral) at the coordinates in millimetres from bregma M/L=±1.5; A/P=-2; D/V=2). Directly afterwards, 100 µl of a preparation of EVs from blood plasma of an LPS injected vaviCre mouse was i.v. injected via the tail vein. Mice were left for 48h to allow for reporter gene expression and were transcardially perfused for further analysis.

### Cage enrichment

Cages used for the experiment were standard cages for mice (365 x 207 x 140 mm, 1284L Eurostandard Type II L). Mice continuously inhabited this cage before an assortment of objects of different sizes and colours were placed in the cage for one hour before they were removed again. Mice were kept for an additional two days before they were killed and brains were removed for further analysis.

### Tissue processing

At the end of the experiment, mice were killed with an overdose of Ketavet i.p. (100 mg/kg)/Xylacin (5 mg/kg) and transcardially perfused with PBS followed by 4% cold PFA in PBS, and brains were removed for subsequent analysis. All conserved organs were post-fixed in 4% PFA in PBS for 24h. For cryosectioning, organs were cryoprotected in 15% sucrose for an additional 24h before they were embedded and serial sectioned (10um) as a frozen block or tissue was snap frozen in methyl butane previously cooled in liquid nitrogen. Sections were attached to glass slides and stored at −20ºC until further use. For fresh fixed tissue, serial coronal sections (30um) were cut on a vibratome (Leica VT1000S) and kept in PBS at 4°C until further use.

### Antibodies

Primary antibodies used in the study were rabbit Calbindin D28-k (Swant, CB-38, 1:5000), rat dopaminergic transporter DAT (Millipore, MAB369, 1:1000), rabbit glial fibrillary acidic protein GFAP (DAKO, ZO334, 1:1000), chicken GFP (Abcam, ab13970, 1:500), rabbit Iba1 (WAKO, 019-19741, 1:1000), mouse NeuN (Millipore, MAB377, 1:300), rat Anti-Mouse CD49d, Clone R1-2 (RUO) (BD Horizon, 1:400), mouse tyrosine hydroxylase TH (Millipore, MAB318, 1:1000), rat anti-mouse CD31 (BD Pharmingen 557355, 1:100) and mouse Olig2 (Millipore MABN50, clone 211F1.1, 1:100).

Secondary antibodies used were Alexa Fluor 647 goat anti-rat (Life Technologies, A21247, 1:1000); Alexa Fluor 488 goat anti-chicken (Abcam, ab150169, 1:1000); Alexa Fluor 568 goat anti-mouse (Invitrogen, A11004, 1:1000); Alexa Fluor 647 goat anti-mouse (Invitrogen, A21235, 1:1000); Alexa Fluor 568 goat anti-rabbit (Invitrogen, A11011, 1:1000) and Alexa Fluor 647 goat anti-rabbit (Invitrogen, A21244, 1:1000).

### Immunohistochemistry

For immunofluorescence staining, brain slices were permeabilized with PBS-Triton X-100 0,1%, blocked in 10% NGS and incubated overnight at 4°C with the primary antibody. On the next day, slices were washed and incubated with the secondary antibody for 2h at 4°C. After a final wash, brain slices were stained with DAPI (Sigma, D9542, 1:1000) and mounted with Aqua-Poly/Mount (18606-20, Polysciences).

For light microscopy, 12µm cryosections were stained on a fully automated Leica Bond III (Leica Biosystems, Germany) using the Leica Bond Polymer Refine Detection Kits (DS9800+ DS9390).

### Image acquisition

Images were captured with either an Epi-fluorescence microscope (Nikon eclipse 80i) or a Confocal inverted microscope (Nikon eclipse TE2000-E). For confocal imaging, a z-stack of pictures of areas of interest was obtained using different picture size magnifications. Images were analyzed with NIS-elements imaging software (version 4.13.05) and ImageJ (https://imagej.nih.gov/ij/). To assess YFP and mCherry expression in the contralateral hemisphere we used a confocal microscope (SP8, Leica, Mannheim, Germany) using multiple objectives at 20x, 40x and 63x objective (HCX PL APO 20x/0,7 dry,HC PL APO 40X/1,40 Oil CS2,HC PL APO 63X/1,40 Oil CS2, Leica). Schematic brain overviews were adapted from http://mouse.brain-map.org/static/atlas.

### Statistical analyses

No statistical methods were used to predetermine sample sizes. Animals were randomly allocated into different experimental groups. No specific randomization method was used. Mice showing incorrect injection sites or optic fiber placement were excluded.

