## Supplemental Figure 1 for "Neuronal activity triggers uptake of hematopoietic extracellular vesicles *in vivo*"

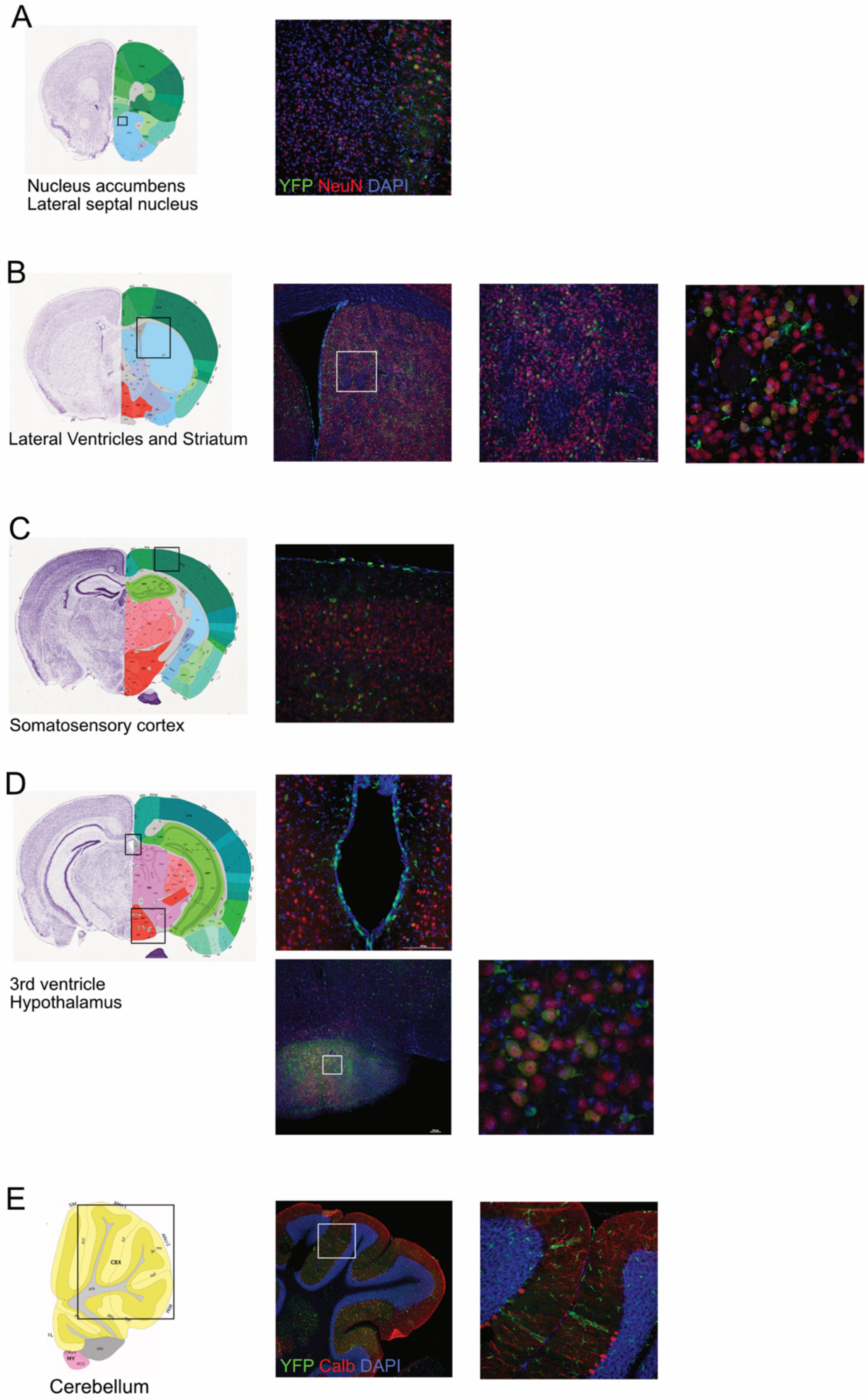


**Supplemental Fig 1. Map of marker gene expression after LPS-induced inflammation in multiple brain areas.**
