## Supplemental Figure 2 for "Neuronal activity triggers uptake of hematopoietic extracellular vesicles *in vivo*"

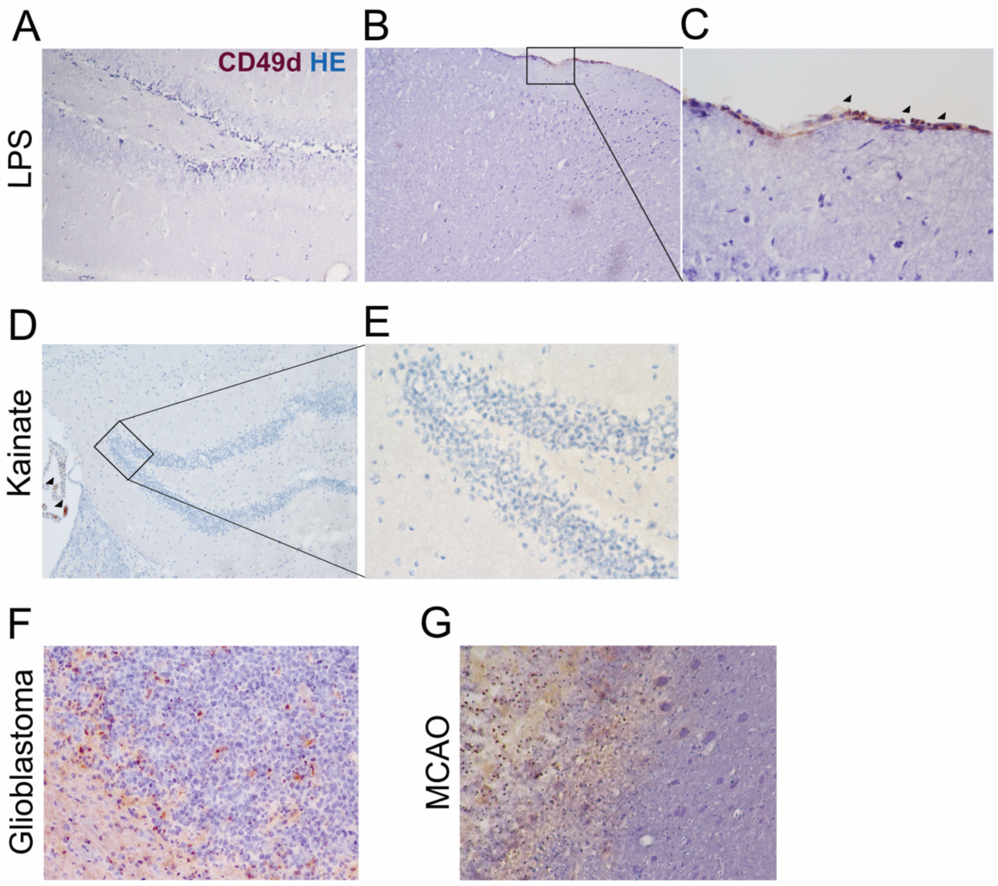


**Supplemental Fig 2. LPS or KA injection does not lead to infiltration of peripheral blood macrophages to the brain parenchyma.** (A) Brain sections from LPS injected mouse do not show CD49d immunoreactive cells in the hippocampus or other brain areas such as cortex (B). (C) Insert with CD49d-positive meningeal macrophage on the brain surface. (D+E) Likewise KA injection does not lead to infiltration of peripheral macrophages. Arrowheads indicate CD49d-positive choroid plexus cells. In conditions causing a high influx of peripheral blood cells into the brain such as glioblastoma (F) and cerebral ischemia caused by MCAO (G) CD49d-positive macrophages are visible.
